# Histone gene expression is regulated by histone H2B ubiquitylation in fission yeast

**DOI:** 10.1101/094565

**Authors:** Viviane Pagé, David Grabowski, C. David Allis, Jason C. Tanny

## Abstract

Cell cycle-regulated expression of histone genes is vital for coordination of DNA replication with chromatin assembly. In yeast, histone genes are regulated by transcriptional activation in S phase and transcriptional repression in other phases of the cell cycle, although the mechanisms are poorly understood. Here we describe a role for histone H2B monoubiquitylation (H2Bub1), a histone mark that is linked to RNA polymerase II transcription elongation, in the activation of replication-dependent histone genes in the fission yeast *Schizosaccharomyces pombe*. Loss of H2Bub1 is also associated with a delay in S phase progression and reduced expression of the replication initiation factor Cdc18 (ortholog of Cdc6). We provide evidence that H2Bub1 impacts histone gene transcription and acts through a mechanism that involves the 3’UTR. Consistent with our previous finding that H2Bub1 is functionally opposed to the elongation factor Cdk9, we also find that the effects of H2Bub1 on histone genes are suppressed by reduction in the activity of Cdk9. Our data suggest that H2Bub1 promotes cell growth by activating cell cycle-regulated genes.

## Introduction

Replication-dependent histone genes are regulated in a cell-cycle dependent manner such that they are upregulated in coordination with S-phase and assembly of replicated DNA into chromatin (1). In metazoans, these genes are distinguished from most other protein coding genes in that they lack introns, and histone mRNA’s lack poly(A)-tails. Instead, these mRNA’s contain a unique stem-loop structure in their 3’UTR that is recognized by SLBP (stem loop binding protein), a specialized RNA-binding protein that mediates histone mRNA 3’ end processing, nuclear export, and translation (2). Metazoan histone genes are also regulated transcriptionally by NPAT (nuclear protein ataxia-telangiectasia locus), which is phosphorylated and activated by cyclin-dependent kinases upon S-phase entry (3).

Unlike their metazoan counterparts, yeast histone mRNA’s are polyadenylated, and are primarily regulated at the level of transcription (4). In the budding yeast *S. cerevisiae*, histone genes are transcriptionally repressed outside of S phase by multiple histone chaperones including HIR (orthologous to the HIRA complex in mammalian cells), Rtt106, and Asf1 (5–7). Activation of transcription in S phase is carried out by a transcription factor complex composed of Spt10 and Spt21, as well as the SBF and MBF transcription factor complexes (8–10). SBF and MBF are dimeric transcription factors composed of a shared regulatory subunit, Swi6, and dedicated DNA-binding subunits (Swi4 in SBF and Mbp1 in MBF). Consensus binding sites for Swi4 and Mbp1 overlap with the Spt10 binding sites upstream of canonical histone genes, suggesting that SBF/MBF and Spt10 function in distinct pathways (11). Transcriptional co-activators for the histone genes include the histone acetyltransferases Gcn5 and Rtt109 and the SWI/SNF chromatin remodeling complex (5,8,12,13).

Less is known about histone gene regulation in the fission yeast *S. pombe.* Involvement of a HIR-type complex in transcriptional repression of histone genes outside of S phase is conserved in *S. pombe* (14). Transcriptional induction of histone mRNA’s in S phase is triggered by binding of the GATA-type transcription factor Ams2 to regulatory elements upstream of histone genes (15). Ams2 expression is activated in G1 by the *S. pombe* MBF complex, composed of the Swi6 ortholog Cdc10 and the DNA-binding proteins Res1 and Res2 (16,17).

Histone H2B monoubiquitylation (H2Bub1) is a universal marker of RNA polymerase II transcription elongation and is an important regulator of cell cycle genes in budding yeast and in mammalian cells (18–21). There is also evidence that H2Bub1 directly promotes DNA replication fork progression and replisome stability in S phase (22). H2Bub1 acts in a histone crosstalk pathway to promote the methylation of histone H3 lysines 4 and 79. H2Bub1 also acts via a methylation-independent pathway whose function is linked to the essential transcription elongation factor Cdk9 (23–25). H2Bub1 and Cdk9 regulate one another through positive feedback mediated by phosphorylation of Spt5, a major Cdk9 target (25). Conversely, genetic evidence indicates that H2Bub1 and Cdk9 have antagonistic functions, although how these are related to expression of individual genes is not known (25). Here we describe a role for co-transcriptional histone H2Bub1 in regulating transcription of replication-dependent histone genes in *S. pombe* that is opposed by Cdk9. Our findings provide insight into the diverse mechanisms through which H2Bub1 regulates cell cycle genes and cell growth.

## Experimental Procedures

### Yeast strains and media

Strains used in this study are listed in Table 1. *S. pombe* strains were cultured in YE (0.5% yeast extract, 3% dextrose) or in EMM (Edinburgh minimal media; (26)) supplemented with adenine, leucine, uracil, and histidine (0.25g/L each). Standard genetic crosses and tetrad dissection was used to introduce the *cdc10-129* allele into other strain backgrounds.

**Table 1.** Strains used in this study.

| Strain number | Genotype | Source |
| --- | --- | --- |
| JTB56 | *h- ade6-M216* | Ref. 24 |
| JTB86-3 | *htb1-K119R::kanMX6 ade6-M216 h+* | Ref. 24 |
| JTB111 | *h- brl2Δ::hphMX4 ade6-M210* | Ref. 24 |
| JTB321 | *cdk9-T212A::kanMX6 leu1-32 ura4-D18 his3-D1 ade6-M210 h+* | Ref. 25 |
| JTB335 | *cdk9-T212A::kanMX6 brl2Δ::hphMX4 leu1-32 ura4-D18 his3? ade6* | Ref. 25 |
| JTB429 | *cdk9-T212A::kanMX6 htb1-K119R::kanMX6 leu1-32 ura4-D18 his3? ade6* | Ref. 25 |
| JTB105 | *htb1::kanMX6 cdc10-129 ade6* | This study |
| JTB106 | *set1Δ::kanMX6 cdc10-129 ade6* | This study |
| JTB107 | *htb1-K119R::kanMX6 cdc10-129 ade6* | This study |

### Synchronization in G1 with ***cdc10-129***

Cells harboring *cdc10-129* were grown to OD_600_ of 0.2 in 300 mL EMM at 25°C and shifted to 36.5°C for 4 hours. Cultures were rapidly cooled to 25°C in ice-water and returned to growth at 25°C in a water-bath shaker. Samples were collected upon return to growth and at 20-minute intervals thereafter. For FACS 1 mL of culture was pelleted in a microfuge, resuspended in ice-cold 70% ethanol, and stored at 4°C. For RNA analyses 10 mL of culture was pelleted, washed with 1 mL sterile dH2O and stored at −80°C.

### RNA analyses

Total RNA was extracted from frozen cell pellets using a hot phenol method (24). For northern blots, 15 μg each sample was electrophoresed on a 1% formaldehyde-agarose gel and transferred to a nylon membrane (Genescreen). Membranes were crosslinked with UV and hybridized to ^32^P-labeled, double stranded DNA probes in 50% formamide, 5X SSC, 5X Denhardt’s solution, 1% SDS, 10% dextran sulfate at 42°C overnight. Probes were generated by PCR with the primers indicated in Table 2, labeled by random priming, purified, denatured, and added to the hybridization mixture (~10^7^ cpm). After hybridization blots were washed with 2X SSC at room temperature, 2X SSC+1% SDS at 65°C, and then 0.1% SSC at room temperature. Blots were exposed to a phosphorimager screen and scanned (Fujifilm FLA-5000).

**Table 2.**
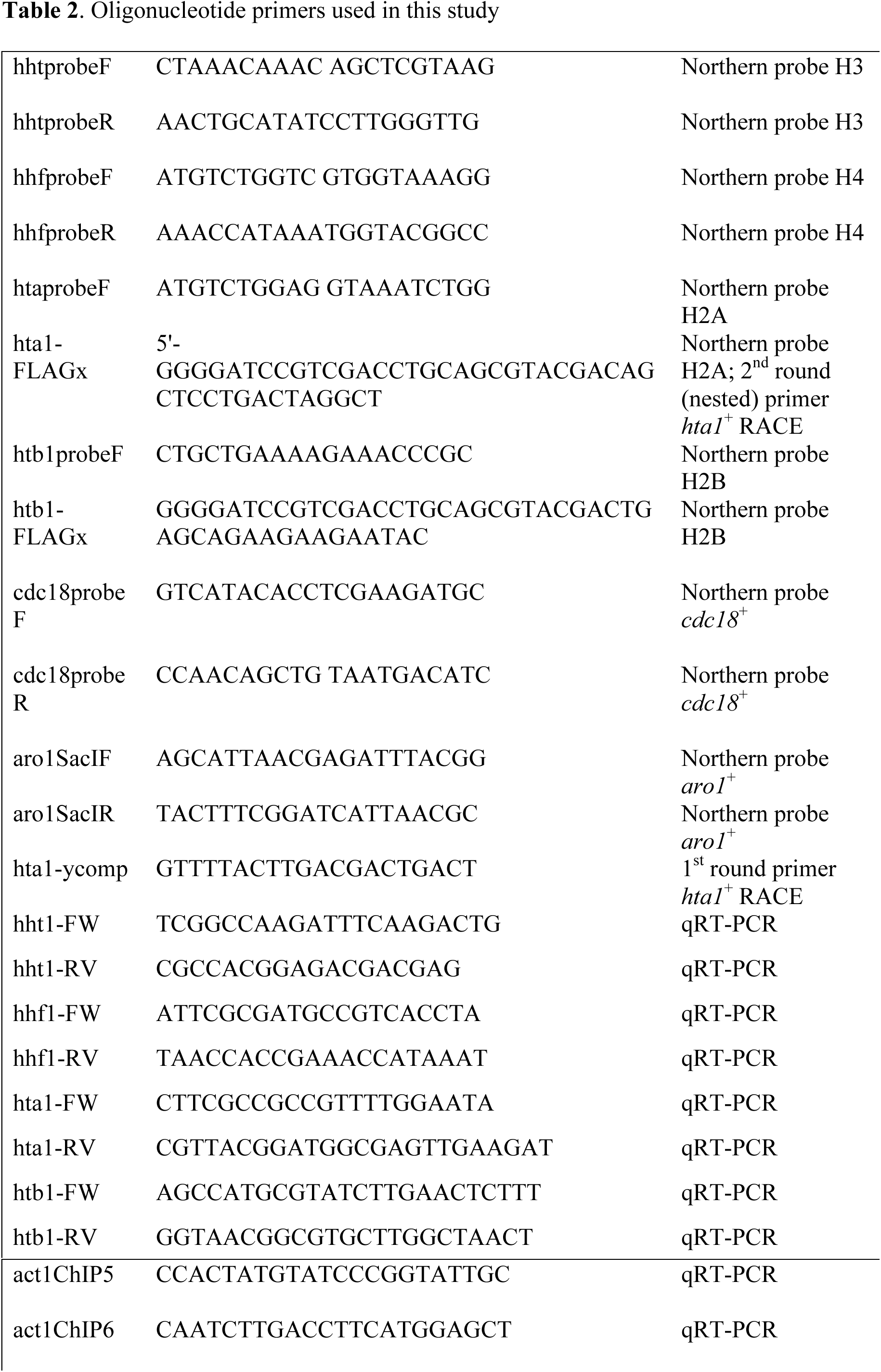
Oligonucleotide primers used in this study.

For RT-qPCR, 1 μg RNA was reverse transcribed to cDNA (Invitrogen Superscript II) with oligo-dT primers, which was then amplified with the primers listed in Table 2 and a SYBR Green qPCR master mix (Bio-Rad) in a Bio-Rad CFX96 qPCR instrument. Levels of histone mRNA’s were expressed relative to *act1*^+^.

5’ and 3’ rapid amplification of cDNA ends (RACE) used total RNA derived from the 100-minute post-release time point for *cdc10-129* and *cdc10-129 htb1-K119R* strains. Analyses were performed using a Generacer kit (Stratagene) and the primers listed in Table 2. Nested PCR (second round) used a 1:100 dilution of the first round PCR reaction as template. Bands corresponding to short cDNAs derived from the *cdc10-129 htb1-K119R* strain were clonded into pCR2.1-TOPO using a TOPO-TA cloning kit (Invitrogen) and sequenced with the M13F primer supplied in the kit.

### Fluorescence-activated cell sorting (FACS)

Samples fixed in 70% ethanol were washed once with 1 mL 50 mM sodium citrate and then resuspended in 0.5 mL 50 mM sodium citrate containing 0.1 μg/mL RNase A. After a 4 hour incubation at 37°C 0.5 mL 50 mM sodium citrate containing 4 μg/mL propidium iodide was added. Samples were sonicated briefly and analyzed on a FACSCalibur-I instrument (BD Biosciences). Data was processed using FloJo software.

## Results

We previously found synthetic lethal interactions between the *htb1-K119R* mutation that removes the consensus ubiquitylation site on histone H2B and deletion of *hip1*^+^ or *slm9*^+^, chromatin assembly factors that repress histone gene expression outside of S phase (24). We thus examined whether *htb1-K119R* caused a similar derepression of histone genes. We isolated RNA from asynchronous wild-type and *htb1-K119R* cells and measured the steady-state levels of mRNA’s encoding all four core histones by RT-qPCR. We found that histone mRNA’s slightly decreased in *htb1-K119R* cells, as well as in cells lacking Brl2, a ubiquitin ligase that catalyzes H2Bub1 (Figure 1). Thus H2Bub1 is a positive regulator of histone gene expression in fission yeast.

**Figure 1.**
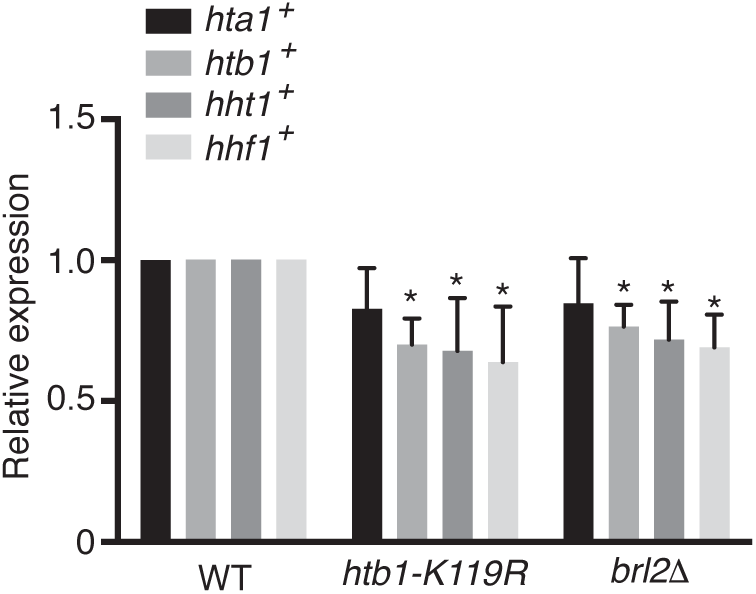
H2Bub1 promotes expression of replication-dependent histone genes in asynchronous *S. pombe* cells. Steady-state RNA levels of the indicated histone genes were quantified by RT-qPCR in the indicated strains (JTB204, JTB86-3, JTB331; see Table 1) and normalized to *act1*^+^ mRNA levels. Wild-type mRNA levels were set to 1. Asterisks indicate significant differences from wild-type (n=3, unpaired t test, p<0.05).

Histone mRNA’s are present at basal levels throughout the cell cycle but are transcriptionally induced in S phase. To determine whether H2Bub1 affects the basal or induced histone mRNA levels, we introduced the temperature-sensitive *cdc10-129* mutation into *htb1-K119R* and control cells to allow synchronization of cells in G1. Both control and *htb1-K119R* cells harbored a *kanMX* cassette (conferring resistance to the antibiotic G418) immediately downstream of the *htb1*^+^ gene; the control genotype will be referred to as *htb1*^+^-*kanMX*. Fluorescence-activated cell sorting (FACS) analysis of control *cdc10-129 htb1*^+^-*kanMX* cells after release from the G1 block revealed a single peak of cells with 1N DNA content. DNA replication, indicated by a transition to a 2N DNA peak, occurred maximally at 100 minutes post-release and was complete by 120 minutes (Figure 2). In *cdc10-129 htb1-K119R* cells arrested at the restrictive temperature, FACS revealed the same 1N peak and a second peak at a position corresponding to >2N DNA content. This second peak is likely the result of cell separation defects caused by the *htb1-K119R* mutation, such that arrest occurs in septated cells harboring two G1 nuclei (24,25). Completion of DNA replication results in formation of a 2N peak and a second peak at a position corresponding to >4N, consistent with persistent failure of cell separation in a fraction of *htb1-K119R* cells. However, the proportion of unseparated cells decreased during the time course, indicating that some separation occurs as cells go through the cell cycle. Importantly, cell cycle progression was delayed in the *htb1-K119R* mutant compared to control, as transition from the 1N to 2N peak was still ongoing at 140 minutes (and to a lesser extent at 180 minutes)(Figure 2).

**Figure 2.**
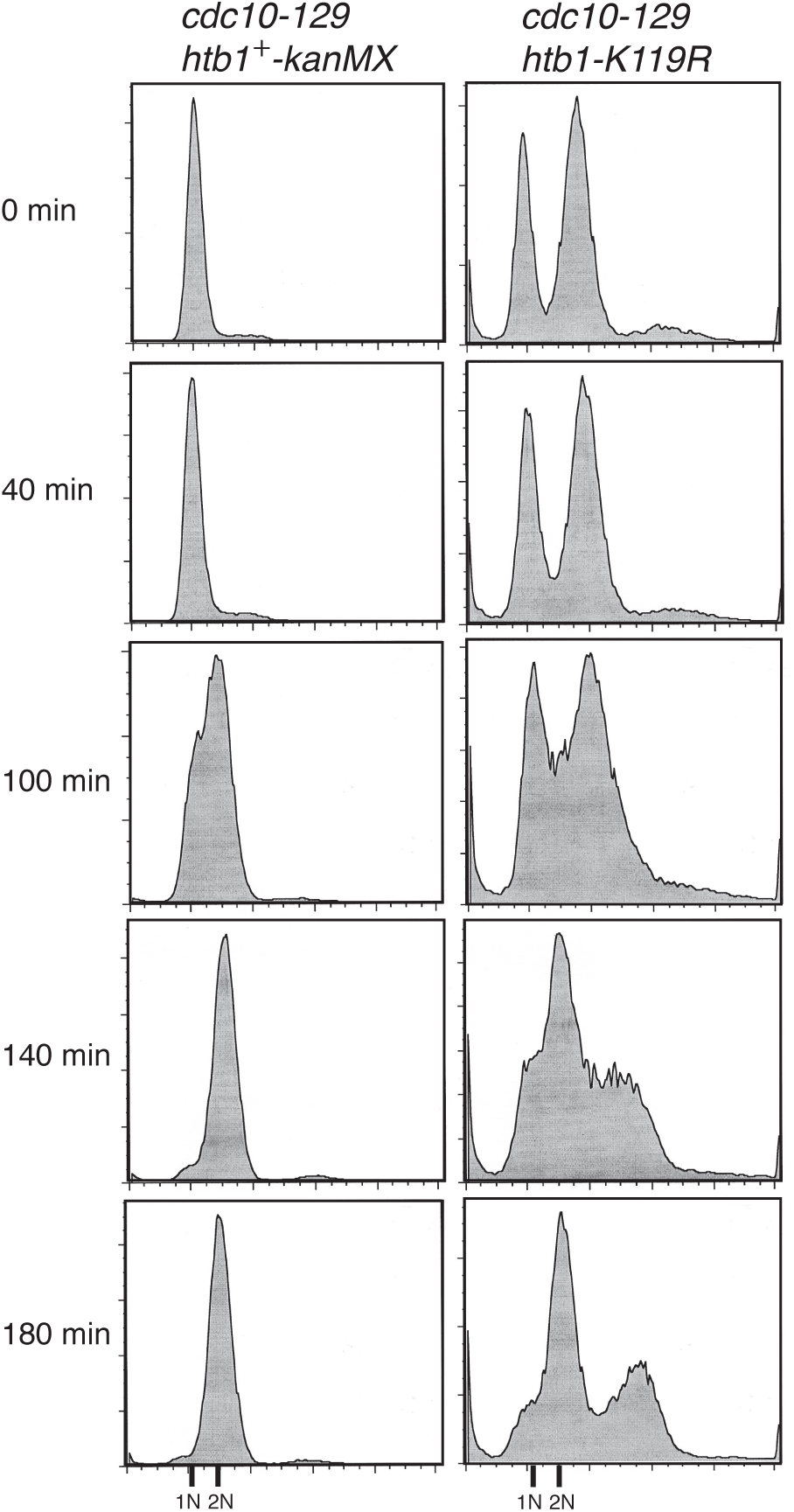
Loss of H2Bub1 delays completion of S phase after release from a G1 block. FACS analysis of *cdc10-129 htb1*^+^-*kanMX* (JTB105) cells or *cdc10-129 htb1-K119R* (JTB107) cells was carried out after synchronization in G1. Times after return to growth at 25°C (in minutes) are indicated on the left. Positions of the 1N and 2N DNA peaks are indicated on the bottom.

We analyzed histone mRNA levels at the same time points by northern blotting (Figure 3A). The *aro1*^+^ transcript, which is not cell-cycle regulated, and total RNA (“EtBr” panel) served as loading controls. In *cdc10-129 htb1*^+^-*kanMX* cells, induction of transcription for genes encoding histones H3, H4, and H2A occurred maximally at 100 minutes post-release, coinciding with the peak of DNA replication (Figure 3). The levels of mRNA’s encoding histones H3, H4, and H2A were markedly decreased in *cdc10-129 htb1-K119R* cells relative to *cdc10-129 htb1*^+^-*kanMX*. This effect was evident throughout the cell cycle. Induction of histone genes still occurred in a manner coincident with DNA replication, albeit at a reduced level. Thus *htb1-K119R* affects levels of histone mRNAs outside of S phase as well as their induction in S phase.

**Figure 3.**
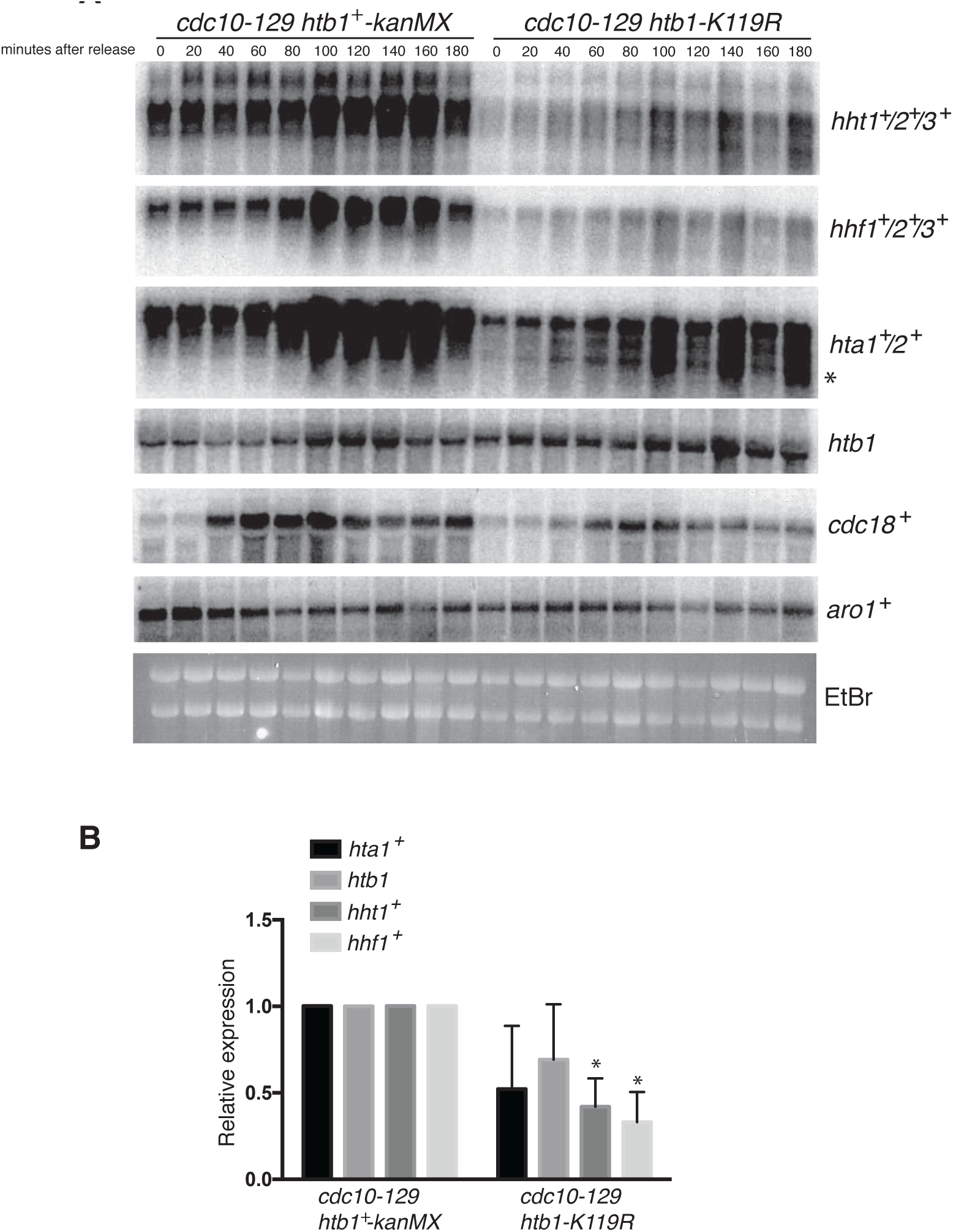
H2Bub1 is required for expression of histone genes and of *cdc18*^+^ after release from a G1 block. (A) Northern analysis of total RNA extracted from cells harvested at the indicated times after release from the G1 block. Probes specific to the indicated genes were used; the *hht, hhf* and *hta* probes each hybridize to all genomic copies of these genes. “*” indicates a short transcript detected specifically in the *cdc10-129 htb1-K119R* strain. “EtBr” indicates the ethidium stain of the agarose gel before transfer; shown are the bands corresponding to 23S and 18S ribosomal RNAs. (B) Quantitative RT-PCR analysis of histone gene expression in the indicated strains (JTB105, JTB107, respectively). Wild-type mRNA levels were set to 1 after normalization to *act1*^+^. Asterisks indicate significant differences from wild-type (n=3, unpaired t test, p<0.05).

Levels of *htb1*^+^-*kanMX* mRNA were lower than those of other replication-dependent histones in the *cdc10-129* background and were less strongly induced in S phase. This was likely due to replacement of the endogenous *htb1*^+^ 3’ UTR with the *kanMX* cassette, a change which by itself does not impact cell growth in asynchronous cells and did not affect kinetics of cell cycle block and release in the *cdc10-129* background (24,27)(Figure 2). Interestingly we found no difference in *htb1* mRNA levels between *cdc10-129 htb1*^+^-*kanMX* and *cdc10-129 htb1-K119R* cells. Since genes encoding all four replication-dependent histones were similarly affected by both *htb1*-*K119R* and *brl2*∆ in asynchronous cells (Figure 1), it is unlikely that this reflects a unique property of *htb1*^+^. Rather, this argues that loss of H2Bub1 affects *htb1*^+^ expression (and perhaps that of other histone genes) through a mechanism that depends on the 3’UTR.

We monitored the expression of *cdc18*^+^, a direct MBF target gene encoding an initiation factor for DNA replication, in our time course experiments in order to verify the cell cycle synchrony (17,28). In *cdc10-129 htb1*^+^-*kanMX* cells *cdc18*^+^ induction after release from the G1 block peaked after 60 minutes. In *cdc10-129 htb1-K119R* cells induction was delayed by approximately 20 minutes, consistent with the delay in DNA replication we observed by FACS. Levels of induction were also reduced, albeit to a lesser extent than what we observed for histone genes. This may point to a more general role for H2Bub1 in cell cycle-regulated gene expression in fission yeast.

We confirmed the histone gene expression defects in the *cdc10-129 htb1-K119R* strain using RT-qPCR. In asynchronous *cdc10-129 htb1-K119R* cells grown at the permissive temperature, *hht1*^+^ and *hhf1*^+^ mRNA levels were significantly reduced relative to those in *cdc10-129 htb1*^+^-*kanMX* cells after normalization to the constitutive *act1*^+^ transcript (Figure 3B). There was no significant difference in *htb1* mRNA levels in these two strains, as observed by northern analysis. The levels of *hta1*^+^ mRNA were also not significantly different by RT-qPCR. We attribute this apparent discrepancy with the northern analysis to the presence of additional *hta1*^+^-derived transcripts in the *cdc10-129 htb1-K119R* strain, as detailed below.

The northern blots revealed the appearance of a short histone H2A transcript in the *cdc10-129 htb1-K119R* strain (Figure 3A; band marked by “*”). Short transcripts are often caused by aberrant transcription initiation associated with defective chromatin structure within gene coding regions (29,30). Using 5’ rapid amplification of cDNA ends (RACE) we mapped the 5’ ends of both full-length and short histone H2A transcripts using primers specific for the *hta1*^+^ isoform (Figure 4A). We mapped the 5’ end of the full-length transcript to a site 24 base pairs downstream of a putative TATA element in the divergent *hta1*^+^-*htb1*^+^ promoter (Figure 4B). The short transcript detected only in the *cdc10-129 htb1-K119R* strain mapped to a site within the *hta1*^+^ coding sequence (Figure 4B). The altered patterns of transcription initiation sites in the *htb1-K119R* mutant support a role for H2Bub1 in transcriptional regulation of *hta1*^+^.

**Figure 4.**
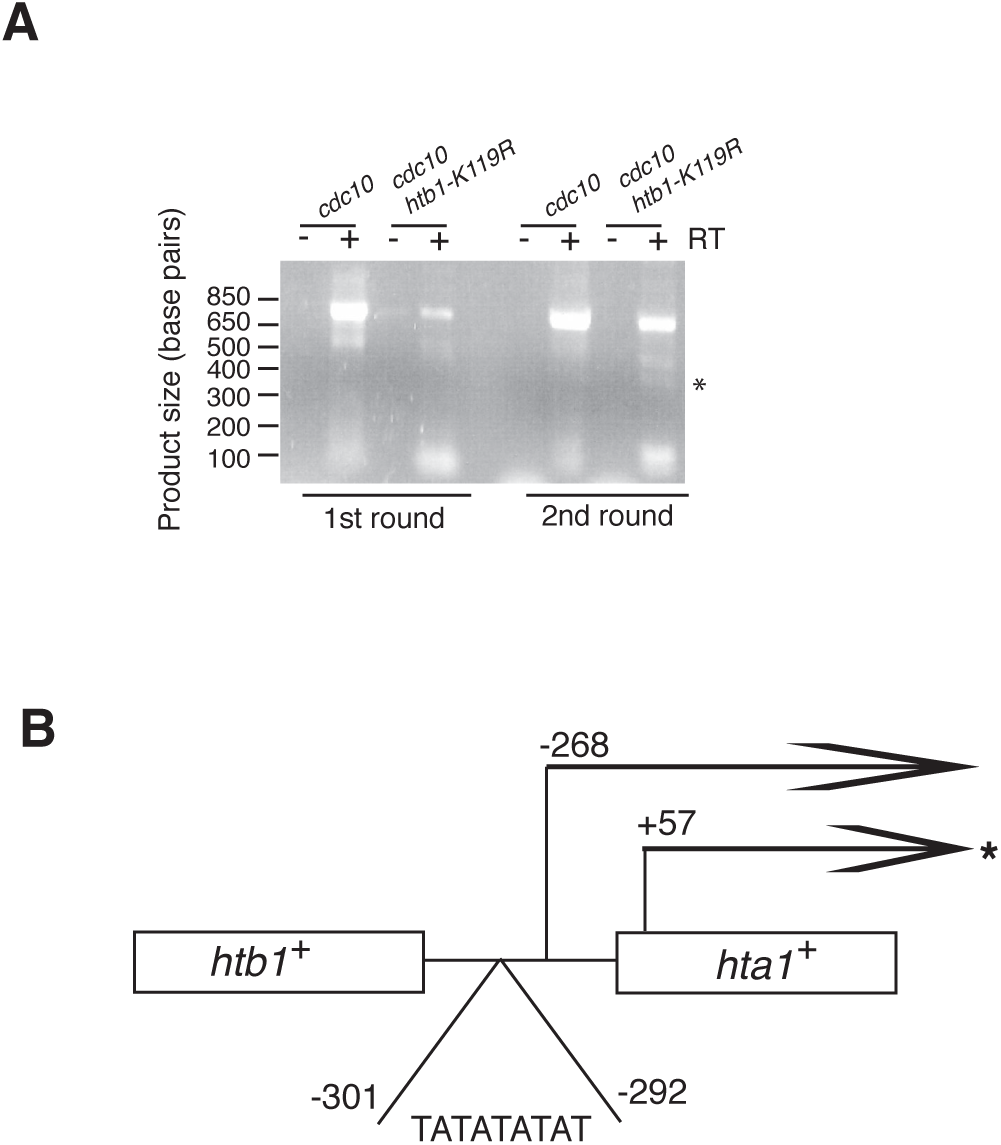
The short *hta* transcript initiates within the coding region of *hta1*^+^. (A) 5’ RACE analysis was performed using the *hta1*^+^ primers listed in Table 2. Shown are PCR products derived from two rounds of amplification of cDNA reactions prepared in the presence (+) or absence (−) of reverse transcriptase (RT). “*” indicates a band derived from a short transcript unique to the *cdc10-129 htb1-K119R* strain. (B) Schematic of the *hta1*^+^-*htb1*^+^ gene pair. Mapped 5’ ends of the full-length and short *hta1*^+^ transcripts relative to the *hta1*^+^ start codon are indicated, as is the putative TATA box.

To determine whether defective histone gene expression was a consequence of loss of H3K4me, we performed a cell cycle block/release experiment on *cdc10-129* cells harboring a deletion of *set1*^+^, encoding the sole H3K4 methyltransferase in S. pombe (31). Histone H2A and *cdc18*^+^ mRNA levels were comparable to those in *cdc10-129 htb1*^+^-*kanMX*, indicating that *htb1-K119R* impacts histone gene expression independently of H3K4me (Figure 5). The robust expression and S phase induction of transcripts produced from the intact *htb1*^+^ gene in the *cdc10-129 set1∆* strain further supported the role of the 3’UTR in maintaining *htb1*^+^ mRNA levels.

**Figure 5.**
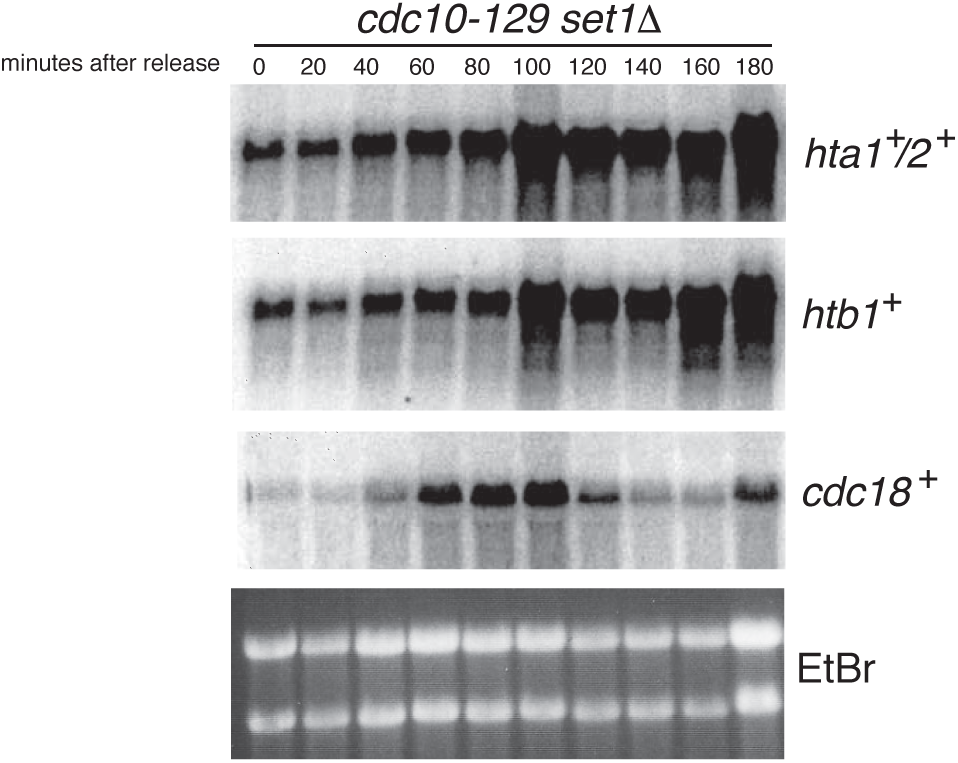
Role of H2Bub1 in histone gene expression is independent of Set1. Northern analysis of total RNA extracted from cells harvested at the indicated times after release from the G1 block. Probes specific to the indicated genes were used. “EtBr” indicates the ethidium stain of the agarose gel before transfer; shown are the bands corresponding to 23S and 18S ribosomal RNAs.

We previously described functional opposition between H2Bub1 and Cdk9, but how this opposition relates to expression of individual genes is unclear (25). To determine whether the effects of htb1-K119R on histone genes might be opposed by Cdk9 activity, we compared *htb1-K119R* cells to double mutant cells that also harbor *cdk9-T212A*, a mutation in the regulatory T-loop that reduces Cdk9 activity (32). The single mutant *cdk9-T212A* cells had wild-type levels of histone mRNA’s during asynchronous growth (Figure 6). Moreover, double mutant cells had histone mRNA levels that were at or above those in the wild-type, indicating that a defect in Cdk9 activity can rescue the histone gene expression defect caused by *htb1-K119R*. Thus Cdk9 and H2Bub1 have opposing effects on histone gene expression in fission yeast.

**Figure 6.**
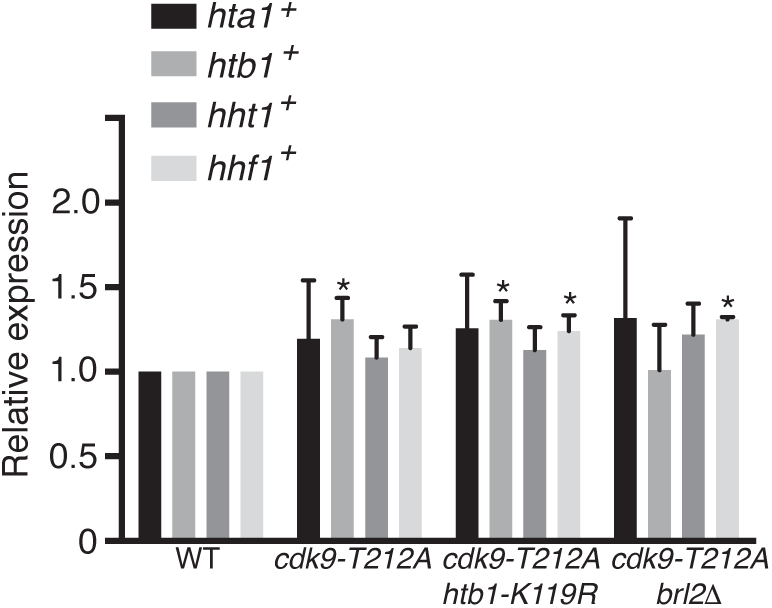
Reduction in Cdk9 activity suppresses the defect in histone gene expression caused by loss of H2Bub1. Steady-state RNA levels of the indicated histone genes were quantified by RT-qPCR in the indicated strains (JTB204, JTB321, JTB429, JTB335; see Table 1) and normalized to *act1*^+^ mRNA levels. Wild-type mRNA levels were set to 1. Asterisks indicate significant differences from wild-type (n=3, unpaired t test, p<0.05).

## Discussion

We have identified a novel role for H2Bub1 in regulation of histone genes in fission yeast. Previous studies in the budding yeast *S. cerevisiae* have linked H2Bub1 to the regulation of cell cycle genes, in that the E2 ubiquitin conjugating enzyme Rad6 and the E3 ubiquitin ligase enzyme Bre1, which target H2B for ubiquitylation, were found to activate cyclin genes in G1 (18). These data point to a role for Rad6/Bre1-dependent H2Bub1 as a co-activator for the SBF transcription factor that drives the G1/S transition. The idea that H2Bub1 and the H2B ubiquitylation machinery might be co-activators for cell cycle transcription factors is supported by findings in human cells, in which the Bre1 ortholog RNF20 has been shown to enhance transactivation by p53 and the androgen receptor (21,33). It is possible that H2Bub1 serves this function for *S. pombe* MBF as well, although histone genes are not direct targets of the orthologous MBF complex and MBF is not limiting for their expression (17). However, our finding that H2Bub1 enhanced activation of *cdc18*^+^, a direct Cdc10/MBF target gene, and the fact that MBF activates Ams2, a transcription factor known to activate histone genes, both suggest that H2Bub1 may have some functional relationship to MBF in *S. pombe*.

H2Bub1 may also act as a co-activator for Ams2. Ams2 is required for the induction of histone mRNA’s in S phase but does not effect their levels outside of S phase, whereas we observe reduction in the levels of histone mRNA’s throughout the cell cycle in the *htb1-K119R* mutant (15). The absence of Ams2 delays completion of DNA replication but appears to have a specific effect on the expression of histone genes (15). The combination of cell cycle genes affected by the loss of H2Bub1 may reflect the importance of this modification in the normal function of multiple transcription factors, as is suggested by the findings in human cells. H2Bub1 may also affect S phase progression via direct effects on the DNA replication machinery, as has been found in budding yeast (22).

The appearance of truncated histone transcripts arising from apparently aberrant transcription initiation within the *hta1*^+^ coding region in the *cdc10-129 htb1-K119R* strain is evidence that loss of H2Bub1 affects histone gene expression at the level of transcription. Intragenic initiation is often indicative of defective transcription elongation, rather than initiation, and is frequently observed when chromatin structure and histone acetylation within gene coding regions is perturbed (29,30,34). H2Bub1 levels correlate with RNAPII elongation rate in gene coding regions (35). Furthermore, there is evidence suggesting that H2Bub1 promotes elongation in concert with the FACT chromatin reorganizing complex, mutations in which lead to accumulation of analogous short transcripts that initiate from within gene coding regions (30,36,37). Although an elongation defect could explain the intragenic initiation phenotype, the pronounced decrease in the levels of the full-length *hta1*^+^ transcript in the *cdc10-129 htb1-K119R* strain suggests that, at this gene, normal transcription initiation is impaired in the absence of H2Bub1. H2Bub1 has been shown to impact transcription of the *ste11*^+^ gene in *S. pombe* through effects at both the promoter and the gene body, although at this gene the effects appear to be opposing (38).

Intriguingly, our data suggest that the defect in histone gene expression may also involve the 3’UTR regions of the histone genes, as the *htb1-K119R* mutation did not affect *htb1* mRNA levels transcribed from the *htb1*^+^-*kanMX* locus that lacks the endogenous 3’UTR. The 3’UTR of the *S. cerevisiae HTB1* gene is important for the cell-cycle periodicity of this transcript, similar to what we observed for *S. pombe htb1*^+^, although the mechanism for this effect is not known (39). Identification of an activity that binds the 3’UTR in *HTB1* mRNA suggests that this element may have a post-transcriptional role (40). However, there is precedent for a transcriptional regulatory function of 3’UTRs. Gene looping, in which the promoter and 3’UTR come into direct physical contact, is common in *S. cerevisiae* and is proposed to promote transcription reinitation and enforce promoter directionality (41,42). Genome-wide chromatin immunoprecipitation experiments have shown that loss of H2Bub1 causes an accumulation of RNAPII at the 3’ ends of genes in *S. pombe*, suggesting that it may preferentially impact transcription of 3’UTRs (25).

Reduction in Cdk9 activity seems to suppress the negative effect of *htb1-K119R* or *brl2∆* on histone gene expression in asynchronous cells, consistent with the phenotypic opposition between Cdk9 and H2Bub1 we have observed previously (25). Cdk9 inactivation may slow elongation sufficiently to normalize transcription through 3’UTR regions in *htb1-K119R* cells, as is suggested by our previous genome-wide ChIP data (25). Alternatively, Cdk9 activity may be required for the repression of histone gene expression outside of S phase, as is suggested by the fact that histone transcript levels are slightly increased in the *cdk9-T212A* mutant strains. In mammalian cells, Cdk9 and H2Bub1 both promote expression of replication-dependent histone genes by enforcing poly(A)-independent transcript termination (43). However, the opposing effects we observe on *S. pombe* histone genes may exemplify how these pathways interact for other poly(A) dependent transcripts. Whether the 3’UTR plays a general role in mediating the effects of H2Bub1 and Cdk9 on gene expression is an important unresolved question.

## Acknowledgements

We thank Paul Nurse for *S. pombe* strains, and members of the Nurse, Allis, and Tanny labs for helpful discussions. This work was supported by the Canadian Institutes for Health Research (M0P-130362 to J.C.T.). J.C.T. is supported by the Fonds de recherche santé Quebec.

